# Protein modelling and thermodynamics reveal that plant cation-chloride cotransporters mediate potassium-chloride symport

**DOI:** 10.1101/2024.05.13.593805

**Authors:** Sam W. Henderson, Saeed Nourmohammadi, Maria Hrmova

## Abstract

Plant cation-chloride-cotransporters (CCC) are proposed to be Na^+^–K^+^–2Cl^−^ transporting proteins, although phylogenetically they are closer to K^+^-Cl^−^ cotransporters (KCC). Conserved features of plant CCC and animal KCC include the presence of predicted K^+^ and Cl^−^ binding sites, and the absence of a Na^+^ binding site. Here, we investigated grapevine (*Vitis vinifera* L.) VvCCC using protein structural modelling and heterologous expression. Our modelling data predicted that 3D folds of VvCCC were more similar to DrNKCC1, but sequences of ion binding sites resembled those of hKCC1. The measurements with *VvCCC*-injected *Xenopus laevis* oocytes with VvCCC, localising to plasma membranes, indicated that oocytes were depleted of intracellular Cl^−^, net ^86^Rb fluxes, which agreed with thermodynamic predictions for KCC co-transport. ^86^Rb uptake by *VvCCC*-injected oocytes was Cl^−^-dependent, did not require external Na^+^, and was partially inhibited by the non-specific CCC blocker bumetanide – these properties are typical of KCC transporters. A loop diuretic-insensitive Na^+^ conductance in *VvCCC*-injected oocytes may account for earlier observations of Na^+^ uptake by plant CCC proteins in oocytes. Our data suggest plant CCC proteins are likely to function as K^+^-Cl^−^ cotransporters, which opens avenues to define their mechanistic and biophysical properties and roles in physiology.

**Highlight:** Based on predictive structural and experimental data, this study presents novel findings to reveal that plant cation-chloride-cotransporters are likely to function as K^+^–Cl^−^ symporters.

## Introduction

Cation-chloride cotransporters (CCCs), classified in the Solute Carrier Family 12 (SLC12) (Zhang *et al*., 2023), mediate electroneutral secondary transport of monovalent Cl^−^ and Na^+^ and/or K^+^ across membranes. Animal cells have three types of CCCs: NKCC proteins that cotransport Na^+^-K^+^-Cl^−^, NCC proteins that cotransport Na^+^-Cl^−^, and KCC proteins that cotransport K^+^-Cl^−^ (Gamba, 2005). Unlike animals, land plants encode fewer CCC proteins in their genomes and plant CCC substrate stoichiometry is unknown (Henderson *et al*., 2018).

Thermodynamics of animal NKCC and KCC were validated experimentally (Russell, 2000). Most animal CCC proteins reside on plasma membranes where ion gradients established by the Na^+^/K^+^-ATPase determine transport direction and physiological function. It was predicted that CCC proteins are accommodated in highly curved sphere-shaped membranes that are bent toward cytoplasmic sides (Lomize *et al*., 2023). Here, KCC proteins mediate the net efflux of K^+^-Cl^−^ down the chemical concentration gradient for K^+^, which is important for inhibitory neurotransmission (Payne, 1997). NCC and NKCC proteins mediate the net influx of Na^+^-K^+^-2Cl^−^ driven by the chemical gradients for Na^+^ and Cl^−^, which is important for solute reabsorption. Conversely, plant CCC proteins reside at the Golgi, *trans*-Golgi network (TGN), and early endosome (EE) membranes (Drakakaki *et al*., 2012; Henderson *et al*., 2015; McKay *et al*., 2022). Their substrates, kinetics and thermodynamics were debated (Teakle and Tyerman, 2010), and even the co-transport of water against its thermodynamic gradient was postulated (Wegner, 2017). However, detailed structural and biophysical studies of plant CCC proteins, which would clarify their transport properties, are lacking.

Plant CCCs are considered to be Na^+^-K^+^-2Cl^−^ symporters (Chen *et al*., 2016; Colmenero-Flores *et al*., 2007; Han *et al*., 2020; Henderson *et al*., 2015; Teakle and Tyerman, 2010). This is because *Xenopus laevis* oocytes expressing AtCCC from Arabidopsis (*Arabidopsis thaliana)* showed greater ^22^Na, ^86^Rb and ^36^Cl radiotracer uptake than water-injected control oocytes (Colmenero-Flores *et al*., 2007); these findings were validated when the orthologous VvCCC protein from grapevine (*Vitis vinifera*) was expressed in oocytes (Henderson *et al*., 2015). Radiotracer fluxes were inhibited by 100 µM bumetanide, which is a loop diuretic and an NKCC antagonist (Colmenero-Flores *et al*., 2007; Henderson *et al*., 2015). However, bumetanide also inhibits KCC proteins, but at higher concentrations (Russell, 2000). Expression of rice (*Oryza sativa*) OsCCC1 in yeast (*Saccharomyces cerevisiae*) led to greater K^+^, Cl^−^, and Na^+^ uptake (Chen *et al*., 2016), further suggesting that plant CCCs may facilitate co-transport of the three ions. Phylogenetically, however, plant CCC proteins reside within or near the KCC rather than NKCC clade across the biological kingdom (Colmenero-Flores *et al*., 2007; Hartmann *et al*., 2014; Henderson *et al*., 2018) (Fig. 1A). Therefore, the present study hypothesised that plant CCCs might display electroneutral KCC (K^+^, Cl^−^) in 1:1 ratio, rather than NKCC (K^+^, Na^+^, Cl^−^) activity.

**Fig. 1:**
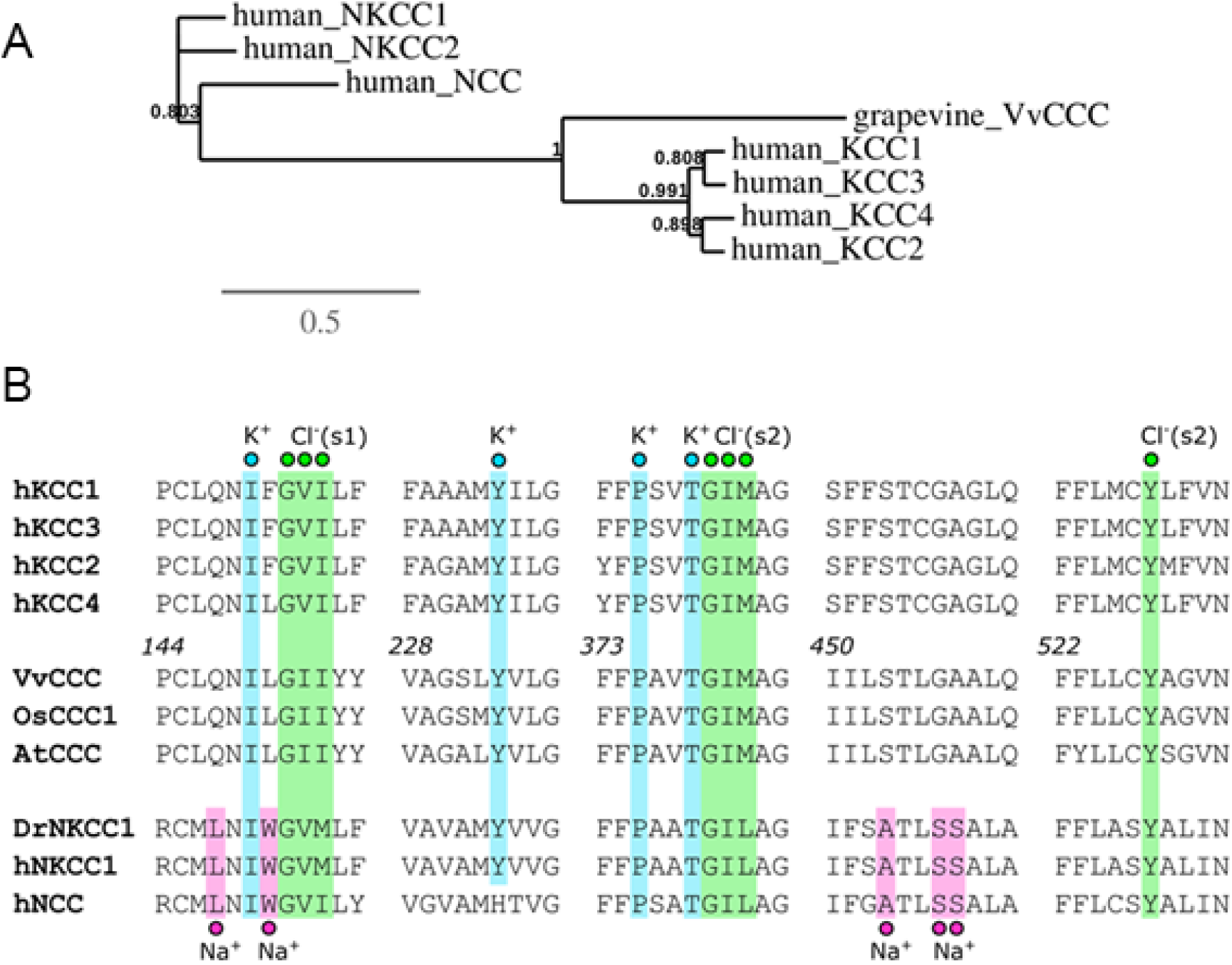
Plant CCC proteins display phylogenetic and structural hallmarks of KCC cotransporters. (A) Maximum likelihood phylogenetic tree of human CCCs and grapevine VvCCC. (B) Amino acid sequence alignment of residues forming the K^+^ (cyan), Na^+^ (magenta), and two Cl^−^ (green) binding sites (s1 and s2) in animal and plant CCC proteins. Numbers indicate the residue positions of the VvCCC protein. Scale = substitution per site.

Cryo-electron microscopy (EM) structures of human NKCC1 (hNKCC1) (Yang *et al*., 2020), zebrafish NKCC (DrNKCC1) (Chew *et al*., 2019), human KCC1 (hKCC1) (Liu *et al*., 2019) and mouse KCC4 (Reid *et al*., 2020) were determined. Based on atomistic simulations it was revealed that human NKCC1 adopts a rocking-bundle mechanism through cooperative angular motions of transmembrane α-helices (Ruiz Munevar *et al*., 2024). These breakthroughs could help elucidate plant CCC function through comparative structural and functional analyses.

The KCC and NKCC proteins form stable homodimers, where monomers form 12 transmembrane α-helical domains (TMD) containing highly conserved residues that form single K^+^ binding sites, and two Cl^−^ binding sites (Delpire and Guo, 2020), although one of the Cl^-^ binding sites (S_Cl1_) in human KCC is believed to be artificial (Liu et *al*., 2019). In addition, NKCC proteins have residues that are capable of coordinating Na ions, and these residues are absent in KCC proteins (Liu *et al*., 2019; Delpire and Guo, 2020). However, critical residues for ion binding in plant CCC proteins remain to be disputed.

This study aims to elucidate the transport properties of VvCCC from grapevine. At the sequence level, VvCCC showed hallmarks of KCC rather than NKCC proteins. The net ^86^Rb flux thermodynamics of VvCCC in *Xenopus* oocytes revealed the KCC-like transport properties. Voltage clamp electrophysiology identified Na^+^ conductance that could explain previous ^22^Na uptake by VvCCC-expressing oocytes. Collectively, these data indicate that VvCCC can function as a K^+^-Cl^−^ cotransporter in heterologous expression systems, which agrees with previous phylogenetic observations. Future work is required to determine precise activity of CCC proteins *in planta*, and whether KCC-like symporter activity is a conserved feature of CCCs across the plant kingdom.

## Materials and methods

### Phylogenetics and sequence alignment

The phylogenetic tree was constructed using the maximum likelihood method implemented in the PhyML v3.0. Amino acid sequence alignment was prepared using ClustalW executed from within Geneious vR9 (Biomatters, New Zealand). Amino acid sequences of the following proteins were used: human hKCC1 (NP_005063.1), hKCC2 (NP_001128243.1), hKCC3 (NP_598408.1), hKCC4 (NP_006589.2), hNKCC1 (NP_001037.1), hNKCC2 (NP_000329.2), zebrafish DrNKCC1 (NP_001002080), hNCC (NP_000330.3), grapevine VvCCC (XP_010655720.1), Arabidopsis AtCCC (NP_849732.1), and rice OsCCC1 (Q6Z0E2.1).

### Identification of the template for homology modelling of Vitis vinifera (VvCCC)

The most suitable template for VvCCC was identified by Phyre2 (Kelley *et al*., 2015) and LOMETS (Zheng *et al*., 2019), and through the evaluations of 21 structures of NKCC and KCC cotransporters retrieved from Protein Data Bank (PDB). These sequences were aligned with ProMals3D (Pei *et al*., 2008) and analysed for distributions of secondary structure elements using PsiPred (Buchan *et al*., 2013).

### 3D Protein molecular modelling

The cryo-EM structures of the cation-chloride cotransporters from *Danio rerio* (PDB accession 6npl, chains A and B) (DrNKCC1) (Chew *et al*., 2019) and human (PDB accession 6m1y, chains A and B) (hKCC3) (Chi *et al*., 2021) in complex with K^+^ and two Cl^−^ ions were used as templates for homology modelling of VvCCC. Both templates represent inward-open inactive states (Chew *et al*., 2019; Liu *et al*., 2019; Zhang *et al*., 2021; Chi *et al*., 2021). In addition, the human hKCC1 (PDB accession 6kkr, chains A and B) (Liu *et al*., 2019) was also used. As the atomic structure of DrNKCC1 does not have defined positions of two Na^+^, in chains A and B, they were docked in TMDs, based on predicted poses (Chew *et al*., 2019; Liu *et al*., 2019; Zhang *et al*., 2021). Top-scoring models were subjected to energy minimisation using the knowledge-based YASARA2 forcefield (bond distances, planarity of peptide bonds, bond angles, Coulomb terms, dihedral angles, and van der Waals forces) (Krieger *et al*., 2002), combined with the particle-mesh-Ewald (PME) energy function for long-range electrostatics (cut-off 8.0 Å) to obtain smoothed electrostatic potentials. To correct covalent geometry, conformational stress was removed by a short steepest descent minimization (time 5000 fs, 1 ft time steps, 298 K), followed by simulated annealing (time step 1 fs, atom velocities scaled down by 0.9 every 10^th^ step) until convergence (710 steps) with energy improvement of less than 0.05 kJ/mol per atom during 200 steps (Krieger *et al*., 2009) in YASARA software. 3D models of VvCCC in complex with three ions (K^+^, and two Cl^−^) using the DrNKCC1 and hKCC3 structural templates were generated in MODELLER 10v1 (Šali and Blundell, 1993) as described (Cotsaftis *et al*., 2012; Waters *et al*., 2013; Xu *et al*., 2018). Best-scoring models from the ensemble of 100 models were selected based on the Modeller Objective Function (Shen and Sali, 2006), Discrete Optimised Protein Energy (Eswar *et al*., 2008), and Statistically Optimized Atomic Potential (Dong *et al*., 2013) terms, PROCHECK (Laskowski *et al*., 1993) and ProSa 2003 (Sippl, 1993). Geometries of pores in TMD domains were calculated by HOLE (Smart *et al*., 1996); these pores are shaped by sphere sequences, where pore radii are equal to their diameters. Structural images were generated in the PyMOL Molecular Graphics System v2.5.2 (Schrődinger LLC, Portland, OR, USA).

The ensembles of 100 structural models of the VvCCC full-length and TMD domain transporters with K^+^ and two Cl^−^ using the DrNKCC1, and hKCC3 templates, were evaluated and top-scoring models were selected for each protein. Stereo-chemical parameters evaluated by PROCHECK (excluding Gly and Pro residues) in DrNKCC1, hKCC3 and VvCCC full-length and TMD structures and models indicated their favourable parameters (Supplementary Table S1), meaning that the VvCCC models were placed in allowed Z-score conformational energy regions (Sippl, 1993), and thus these models were reliable.

### Structural bioinformatics

Sequence conservation patterns of VvCCC, DrNKCC1 and hKCC3 were analysed with ConSurf (Celniker *et al*., 2013; Landau *et al*., 2005) at sequence identities between 35% and 95% (specifications: HMMMER homolog search algorithm, UNIREF-90 Protein database with the E-value cut off of 1·10^-4^, Bayesian Model of substitution) based on the top-scoring VvCCC models.

### Thermodynamic predictions

Thermodynamic driving forces for KCC proteins with K^+^:Cl^−^ stoichiometry (ΔμKCC) were calculated at different [Cl^−^]_i_ using: 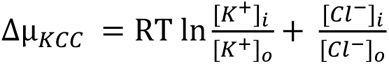 assuming 100 mM [K^+^]_i_. Thermodynamic driving forces for NKCC proteins with Na^+^:K^+^:2Cl^−^ stoichiometry (ΔμNKCC) were calculated using: 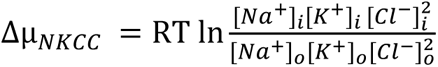 assuming 100 mM [K^+^]_i_ and 10 mM [Na^+^]_i_. Assumed intracellular Na^+^ and K^+^ concentrations fall within the typical range reported for *Xenopus laevis* oocytes (Sobczak *et al*., 2010). Extracellular Cl^−^ was set to 106.6 mM [Cl^−^]_o,_ which is the concentration within ND96 solution. Subscripts i and o denote intracellular and extracellular compartments, respectively, R is the gas constant (8.314 J mol^−1^ K^−1^) and T is the absolute temperature (298.15 °K).

### Xenopus oocyte experiments

Radiotracer flux experiments were performed as described (Henderson *et al*., 2015). Briefly, VvCCC cRNA was sythesized using the T7 mMessage mMachine kit (Thermo Fisher Scientific Inc., MA, USA). Oocytes were injected with 25 ng of cRNA and incubated in Ringer’s solution at 18 °C for two days. Oocytes were pre-incubated in a Cl^−^-free ND96 solution (Cl^−^ replaced with gluconate) for 2 hours to overnight, to deplete internal [Cl^−^]_i_. Oocytes were transferred to flux solutions containing 1 µCi of ^86^Rb radionuclide (as RbCl) (NEZ072001MC, Perkin Elmer). Flux solutions consisted of standard ND96 (96 mM NaCl, 2 mM KCl, 1.8 mM CaCl_2_, 1 mM MgCl_2_, 5 mM HEPES, pH 7.4), or ND96 where 2 mM KCl was replaced with 10 mM KCl or standard ND96 where 2 mM KCl was replaced with 5 mM RbCl. In both cases, the NaCl concentration was adjusted to maintain osmolality. A flux solution was prepared consisting of ND96 where 96 mM NaCl was replaced with NMDG-Cl. All flux solutions contained 0.1 mM ouabain to block the oocyte Na^+^/K^+^-ATPase, and some solutions included 100 µM bumetanide. Oocytes were preincubated in inhibitors for 10 minutes prior to being transferred to a flux solution. Oocytes were rinsed three times in an ice-cold flux solution without radionuclide, transferred individually to scintillation vials, and lysed with 10% (w/v) SDS. The scintillation cocktail was added and samples were counted using a liquid scintillation counter (Beckman LS6500). Counts per minute were converted to pmoles per oocyte using: 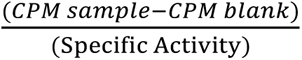. The Specific activity (SA) of 10 µl aliquots of flux solution was calculated according to:

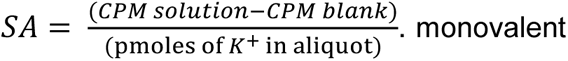

Two-electrode voltage clamping was performed in a standard isotonic Ca^2+^-free ND96 solution (100 mM NaCl, 2 mM KCl, 5 mM MgCl_2_, 10 mM HEPES, pH 7.4). Resting membrane potentials (V_m_) and whole-cell currents were recorded using a GeneClamp 500B amplifier (Molecular Devices LLC., CA, USA) connected to two microelectrodes made of borosilicate glass that were backfilled with 1 M KCl. From a holding potential of −40 mV, oocytes were clamped stepwise from −120 mV to +60 mV for 300 ms and whole-cell currents were recorded. Data were digitized using a Digidata 1322A (Molecular Devices LLC.), and data were analysed using pClamp version 9.0 software (Molecular Devices LLC.).

### Microscopy

An expression construct was assembled to visualise the subcellular localisation of VvCCC-DsRed in HEK293 cells. pcDNA3.2-DEST vector backbone (Thermo Fisher Scientific) was linearised by PCR using primers 5’-TTGATCTAGAGGGCCCGCG-3’ and 5’-TGATAGCTTAACTAGCCAGCTTGG-3’. Full-length VvCCC coding sequence without stop codon was amplified from pCR8-VvCCC (Henderson *et al*., 2015) using primers 5’-CTAGTTAAGCTATCACACCATGGACAACGGAGACATTGA-3’ and 5’-CGATCCTCCTCCTCCTGTGAAAAGGGTGACAACATCT-3’. DsRed was amplified from pAG423GAL-ccdB-DsRed (Addgene plasmid repository, plasmid #14365) using primers 5’-GGAGGAGGAGGATCGATGGACAACACCGAGGACGT-3’ and 5’-GGCCCTCTAGATCAACTACTGGGAGCCGGAGTGG-3’. PCR was performed using Platinum™ PCR SuperMix High Fidelity (Thermo Fisher Scientific) containing 0.8 µM of forward and reverse primer, plus 2.5 ng of a plasmid template. Purified PCR fragments were assembled using Gibson Assembly® Master Mix (New England Biolabs, MA, USA) following the manufacturer’s procedures.

Human Embryonic Kidney 293 (HEK293) cells were grown at 37 °C with 5% (v/v) CO_2_ in Dulbecco’s modified eagles medium (DMEM). Cells were cultured to 80% confluence in an ibidi 8-well µ-slide (Ibidi GmbH, Germany) and transfected with 1 µg of pcDNA3.2-DEST-VvCCC-DsRed using lipofectamine 3000 reagent (Thermo Fisher Scientific). After two days, the cells were stained for 15 minutes at 37 °C with Hoechst nuclear stain and CellMask™ Green Plasma Membrane Stain (Thermo Fisher Scientific). Cells were washed 3 times with phosphate-buffered saline to remove an unbound stain, resuspended in DMEM and imaged using an FV3000 Confocal Microscope equipped with an x40 oil immersion objective lens (Olympus Australia Pty Ltd, Australia). Excitation/emission conditions were Hoechst (405 nm/430–470 nm), CellMask™ (488 nm/500–540 nm) and DsRed (561 nm/570–670 nm).

For yeast, VvCCC-YFP was cloned into pYES-DEST52 (Thermo Fisher Scientific) by LR recombination with pCR8-VvCCC-YFP (Henderson *et al*., 2015) using Gateway LR clonase (Thermo Fisher Scientific). The expression construct was introduced into *Saccharomyces cerevisiae* strain B31 (MATα ena1–4::HIS3 nha1::LEU2) (BaAueIos *et al*., 1998) using the lithium acetate procedure. Transformants were selected on a synthetic complete medium without uracil (SC-uracil) at 28 °C. Yeast cells were grown overnight in liquid SC-uracil containing 2% (w/v) ᴅ-glucose at 28 °C with shaking. To induce protein expression, cells were pelleted and resuspended in SC-uracil containing 2% (w/v) ᴅ-galactose and incubated overnight at 28 °C with shaking. Prior to imaging, cells were incubated in a growth medium containing 10 µg/mL of FM4-64 (Sigma, St Louis, MO, USA) which stains the plasma membrane of living cells before being internalised (Vida and Emr, 1995). Cells were rinsed and imaged using a Nikon A1R confocal laser scanning microscope with ×63 water objective lens and NIS-Elements C software (Nikon Corporation, Tokyo, Japan). Excitation/emission conditions were GFP (488 nm/500–550 nm) and FM4-64 (561 nm/570–620 nm).

## Results and Discussion

### Plant and animal CCC proteins share residues for K^+^ but not for Na^+^ binding

Cryo-EM structures of animal CCCs revealed critical residues for ion binding. NKCC and KCC monomers contain single K^+^ binding sites where the cation is coordinated by five residues at distances of around 3 Å (Chew *et al*., 2019; Liu *et al*., 2019). Alignment of three canonical plant CCCs (VvCCC, AtCCC and OsCCC1) with structurally resolved animal CCCs showed that the residues forming the K^+^ binding sites were conserved (Fig. 1B). Animal CCC monomers contain two Cl^−^ binding sites where ions are coordinated by backbone amide groups of three consecutive residues, plus a fourth residue at the second site (Chew *et al*., 2019; Liu *et al*., 2019). In plants and animals, the residues form similar signatures, when in the first putative Cl^−^ site (Cl^−^) aliphatic isoleucine replaces aliphatic valine at an equivalent position 151 of VvCCC/AtCCC/OsCCC1 (Fig. 1B). The amide group of isoleucine in plant CCCs could form an anion binding site, as this is the case for many anion receptors (Gale *et al*., 2016). The second Cl^−^ site is significantly conserved between aligned plant and animal CCCs (Fig. 1B). The NCC and NKCC proteins also harbour A535, S538 and S539 (in *Danio rerio* NKCC1) that form Na^+^ coordination sites (Chew *et al*., 2019; Yang *et al*., 2020). Based upon the primary sequence analysis, plant CCC proteins, like human KCCs, do not share those Na^+^-binding residues (Fig. 1B), indicating that VvCCC may coordinate neutral K^+^-Cl^−^ ion pair rather than Na^+^-K^+^-2Cl^−^ for co-transport.

### Structural modelling reveals that K^+^ and not Na^+^ is the preferred cation for VvCCC

Structurally modelling was performed to further investigate the ion binding of plant CCC proteins. The most suitable template for VvCCC identified by Phyre2 and LOMETS was the DrNKCC1 cryo-EM structure in complex with K^+^ and two Cl^−^ ions (Chew *et al*., 2019). The second and third considered templates were hKCC1 (Liu *et al*., 2019) and hKCC3 (Chi *et al*., 2021), both in complex with K^+^ and two Cl^−^ ions. In these structures, the two Cl^−^ ions bind at Cl^−^_1_ and Cl^−^_2_ sites, where the second binding site is located at the surface of the structure, and is considered to be artificial; but the possibility remains that it could also support the structural integrity of K^+^ binding sites in animal KCCs (Chi *et al*., 2021; Liu *et al*., 2019). hKCC1 and hKCC3 templates were considered less favourable for modelling due to substantial mismatches in secondary structure distributions between TMDs of VvCCC and both hKCCs, in addition to a significantly higher gap presence (Supplementary Table S1). These secondary structure mismatches and gaps resulted from the presence of a 107-residue loop in the TMD domain (between α-helices 5 and 6) of hKCC1, which is shorter in hNKCC1 (26 residues) and VvCCC (29 residues). hKCC1 contains a large extracellular 120-residue linker between transmembrane helix (TM) 5 and TM6a, and this domain is unique to the KCC subfamily in animals and likely forms a gate (Delpire and Guo, 2020). It is therefore possible that plant and animal CCC proteins are differentially gated, although further research is required to confirm this.

Further comparisons of TMD displacement parameters indicated a higher structural similarity in packing between DrNKCC1 and VvCCC (or AtCCC1) models with the DrNKCC1 template compared to any of the KCC templates. This further validated that DrNKCC1 served as a more suitable template for 3D modelling of VvCCC (Supplementary Fig. S1).

Three dimensional models of full-length VvCCC revealed the overall architecture consists of a dimeric quaternary assembly (each dimer with two protomers) related by a pseudo-twofold symmetry axis. These folds formed the N-terminal α-helical TMD bundles and C-terminal mixed CTD α/β folds (Fig. 2). K^+^ and two Cl^−^ ions were bound in pores of TMD domains (Fig. 2; Fig. 3). The docked Na^+^ ion within DrNKCC1 was placed in a small cavity, bounded by α-helical breaks and around four short α-helices, at around 6-7 Å distance from the K^+^ ion in both dimers (Fig. 2). No major structural differences in 3D models between TMD domains of VvCCC with three (K^+^, and two Cl^−^) or four (K^+^, Na^+^ and two Cl^−^) ions were observed. It would appear that both cations could compete for binding sites in pores and cavities with opposing strengths (Chew *et al*., 2019; Chi *et al*., 2021; Liu *et al*., 2019; Zhang *et al*., 2021). The AlphaFold Protein Structure Database (Jumper et *al.*, 2021) contains the 3D monomeric VvCCC model (accession AF-F6HLW8; likely modelled on human KCC1), with the root-mean-square-deviation (rmsd) value of 1.8 Å. We also generated the dimeric model of VvCCC using Cosmic^2^ (Cianfrocco *et al*., 2017) with the rmsd value of ∼6 Å to VvCCC (modelled on DrNKCC1) and nearly identical distributions of secondary structure elements.

**Fig. 2:**
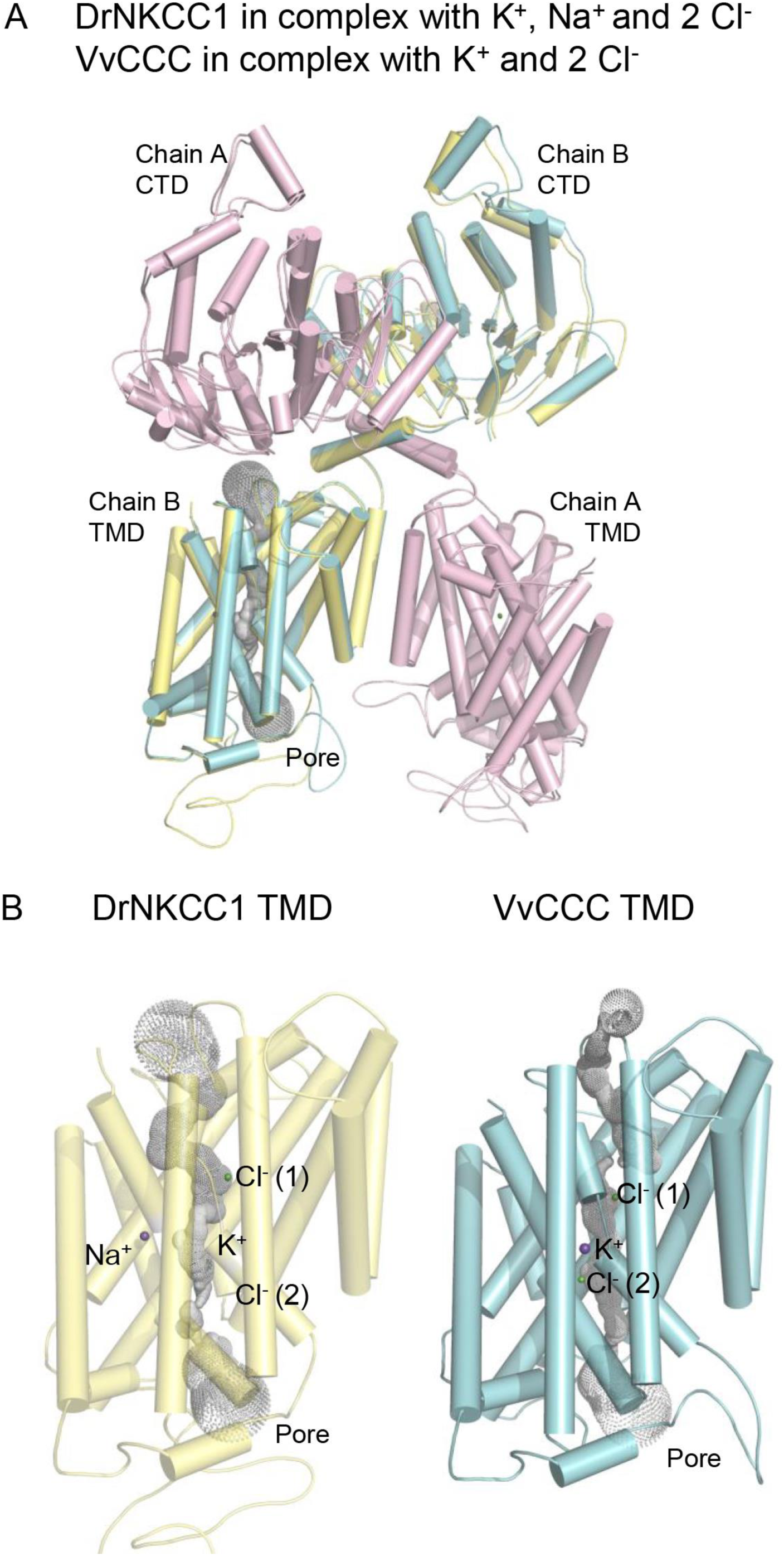
Cryo-EM structure of DrNKCC1 and full-length molecular model of VvCCC in complex with ions. (A) Cartoon representations of superposed DrNKCC1 and VvCCC viewed from the membrane plane (rmsd value 0.94 Å over Cα carbons of 1,720 and 1,696 residues, respectively). Cartoons illustrate spatial dispositions of dimeric assemblies with cylindrical α-helices in chains A (pink for DrNKCC1, yellow and cyan for VvCCC), with chains A and B forming pores (calculated by Hole; Smart *et al*., 1996), shown in dots in chains B. Spatial dispositions of TMDs (α-helical bundles) relative to CTDs (mixed α/β fold domains; chains A and B) are indicated. The pore of DrNKCC1 accommodates K^+^ and two Cl^−^ ions (cpk spheres) with Na^+^ bound nearby, while pores of VvCCC accommodate K^+^ and two Cl^−^ ions. (B) Cartoon representations of TMD domains (chains B) of DrNKCC1 (yellow), and VvCCC (cyan), illustrating the poses of K^+^ and two Cl^−^ ions (cpk spheres) contained in pores.

**Fig. 3:**
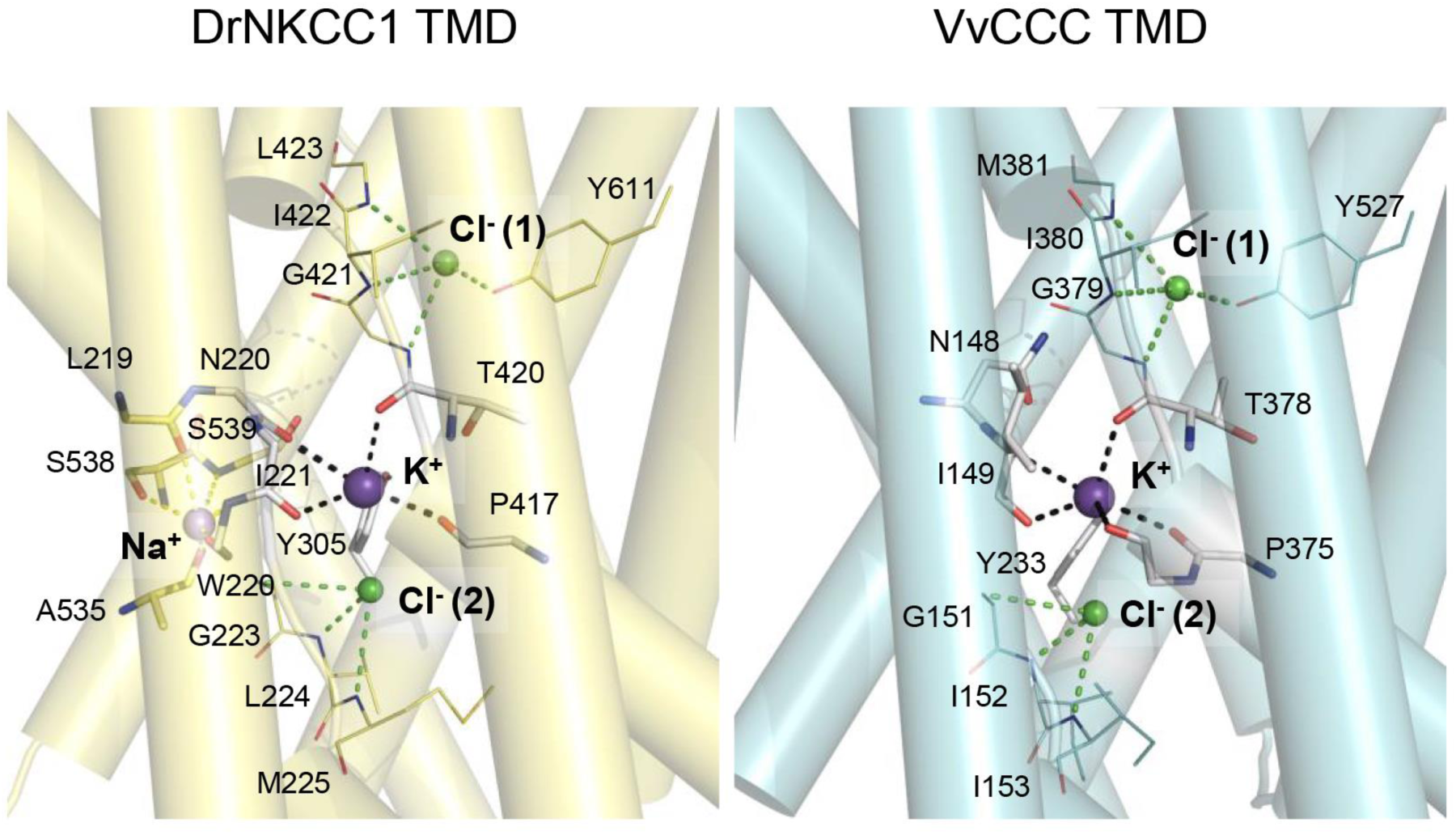
Close views of the cryo-EM structure of DrNKCC1 and the molecular model of VvCCC in complex with ions. Detailed views of TMD domains (chains B) of DrNKCC1 (left panel), and VvCCC (right panel) illustrating the poses of K^+^, (also Na^+^ in DrNKCC1) and two Cl^−^ ions [Cl^-^ (1) and Cl^-^ (2)] (cpk spheres) in pores. Residues participating in ion binding are indicated: yellow cpk sticks for Na^+^-binding, cpk sticks for K^+^ -binding, and yellow lines for two Cl^−^-binding residues in DrNKCC1, and cpk sticks and cyan lines for residues that bind K^+^ and two Cl^−^ in VvCCC. Separations in black dashed lines between residues and K^+^, Na^+^ and Cl^−^, are indicated. These separations are between 2.8 Å and 3.1 Å (K^+^), 2.1 Å and 3.0 Å (Na^+^), and 3.1 Å and 3.7 Å (two Cl^-^) for DrNKCC1, and between 2.8 Å and 3.6 Å (K^+^), and 3.3 Å and 3.8 Å (two Cl^-^) for VvCCC.

ConSurf and ProMals3D sequence analyses of residues that participated in the binding of K^+^, Na^+^ and Cl^−^ ions showed that residues at the K^+^, and two Cl^−^ binding sites were highly conserved between DrNKCC1, hKCC1, and VvCCC (Fig. 3). Conversely, the sites for the second cation (Na^+^ or K^+^) showed weak conservation between VvCCC and DrNKCC1, but strong conservation between VvCCC and hKCC1 (Fig. 3). These comparisons suggested that although VvCCC and DrNKCC1 were structurally similar, the Na^+^ binding site in VvCCC resembled that of hKCC1 (Fig. 3). The other possibility coexists that VvCCC could bind two K^+^ and two Cl^−^ ions, where the pose of the second K^+^ ion would be in the predicted Na^+^ pose of DrNKCC1 (Chew *et al*., 2019; Liu *et al*., 2019; Zhang *et al*., 2021). Ion binding sites showed characteristics consistent with cation or anion environments, where in the former case, carbonyl oxygens participated in the binding of cations, while main-chain amide groups secured the poses of anions. The cation binding sites of cotransporters were reminiscent of Na^+^ binding sites in HKT transporters, where carbonyl oxygens participated (Cotsaftis *et al*., 2012; Houston *et al*., 2020; Waters *et al*., 2013; Wege *et al*., 2021; Xu *et al*., 2018).

ConSurf sequence conservation patterns (Table 1), based on the top-scoring model of VvCCC and AtCCC in complex with ions, revealed high conservation scores (9,9 and 9,8) of residues involved in K^+^ binding, and those of Cl^−^ and Cl^−^ sites (9,8 to 9,9), although some variations in ion binding residues occurred. The two Ser453 and Gly456 residues that could participate in the binding of an additional cation in VvCCC (Table 1), were also conserved amongst 150 homologous sequences, except Ala457, which showed a lower conservation pattern and higher variability (Table 1). It could be predicted from the structural analyses that VvCCC and AtCCC1 bind K^+^ and Cl^-^ in 1:1 ratio or K^+^ and Cl^-^ in 2:2 ratio to assure electroneutral binding. The third possibility to bind K^+^, Na^+^ and Cl^-^ in 1:1:2 ratio is less likely given the conservation patterns of a putative Na^+^ (or other cation) binding site (Table 1). This is supported by ConSurf analyses of 150 homologous sequences to VvCCC and 250 sequences to AtCCC, where the STL(C)GA signature for the second cation binding (Table 1; Fig. 1b; Fig. 3) is absolutely conserved between VvCCC or AtCCC and hKCC1 – thus, this analysis suggests that Na^+^ binding is unlikely.

**Table 1:**
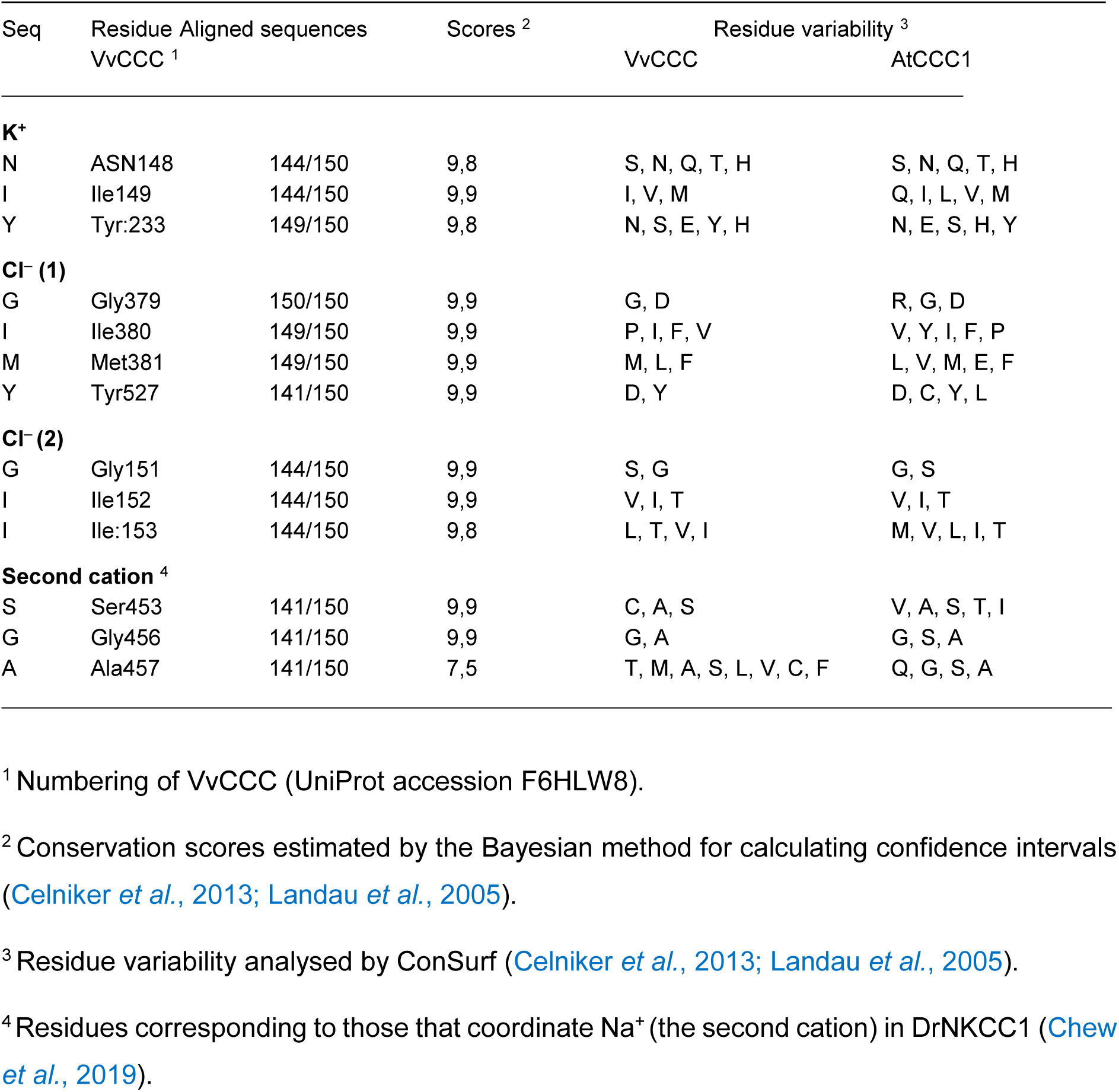
Residue conservation scores and patterns involved in the binding of K^+^ and two Cl^−^ ions and a putative second cation by VvCCC and AtCCC1, at sequence identities between 35% and 95%, analysed by ConSurf.

In summary, although 3D modelling demonstrated that VvCCC shared structural features closely related to animal KCC proteins (*e.g.* DrNKCC1), the dispositions of ion binding residues indicated that VvCCC is capable of K^+^ binding and incapable of Na^+^ ion binding.

### VvCCC is intracellular in yeast and HEK293 cells

Transport properties of CCCs have traditionally been investigated in *Xenopus* oocytes using radiotracers such as ^86^Rb, which by its atomic nature mimics Na^+^ and K^+^ alkali metals. However, radionuclides are hazardous and becoming more difficult to source. So, alternative methods using fluorescent probes in mammalian cells, or whole-cell perforated patch-clamping, are often used (Medina and Pisella, 2020). These assays require that CCCs localise to the plasma membrane. In HEK293 cells, a VvCCC-DsRed fusion protein showed an intracellular signal that did not co-localise with plasma membrane (CellMask^TM^) or nuclear (Hoechst) stains (Fig. 4A). Hence, HEK293 cells could not be used for functional transport studies. Factors influencing VvCCC subcellular localisation and its precise membrane targeting in HEK293 cells remain to be determined, although it is known that phosphorylation could influence the plasma membrane targeting of hKCC2 in HEK293 cells (Lee *et al*., 2010; Lee *et al*., 2007).

**Fig. 4:**
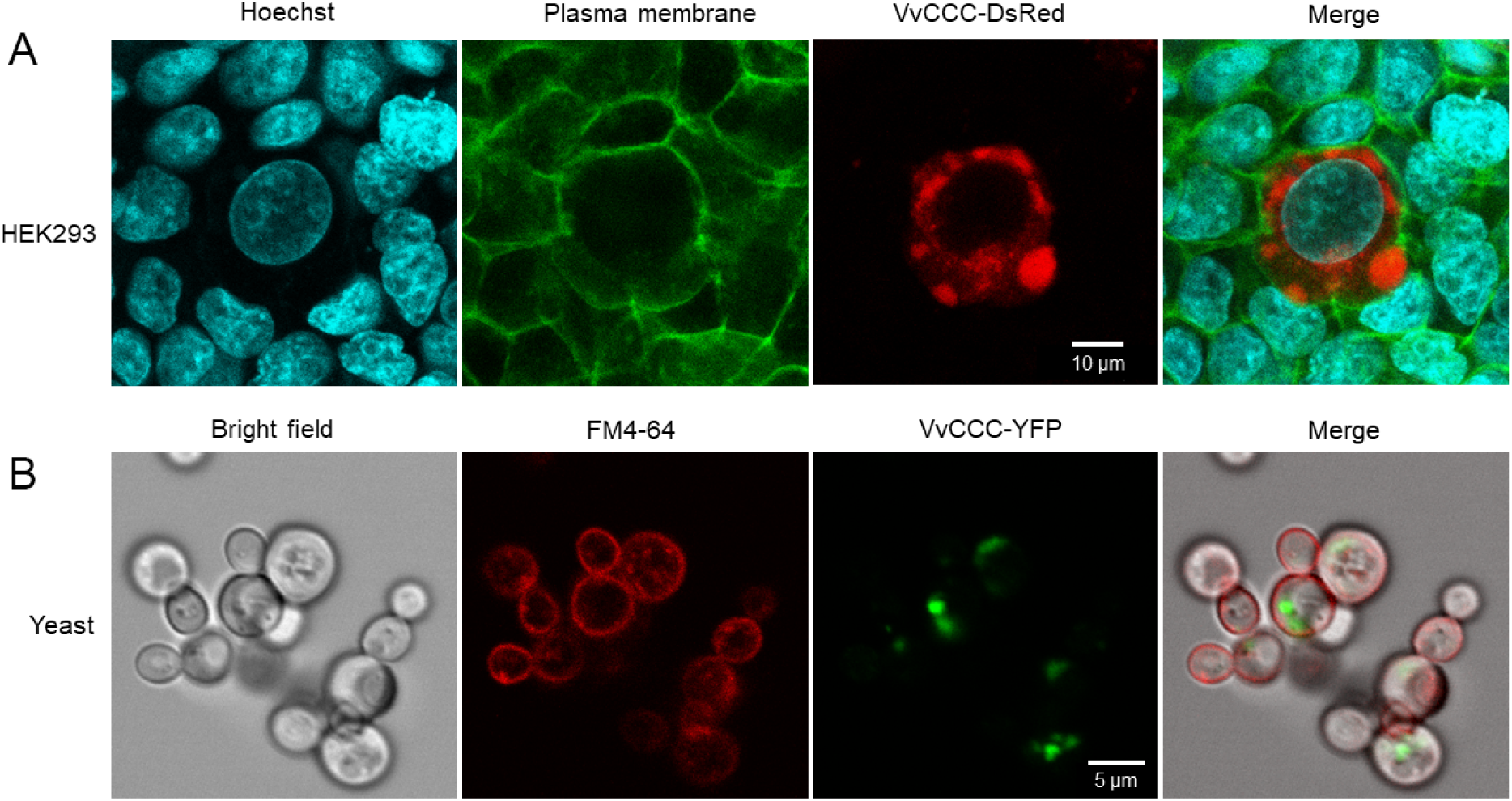
Fluorescently tagged VvCCC localises to intracellular compartments of yeast and HEK293 cells. (A) Transient expression of VvCCC-RFP in transfected HEK293 cells. Cells were counter stained with Hoechst nuclear dye and CellMask^TM^ plasma membrane stain and imaged by confocal microscopy. (B) Transient expression of VvCCC-YFP in yeast (*Saccharomyces cerevisiae*). Cells were counter stained with FM4-64 as a membrane marker and imaged by confocal microscopy.

Growth of Na^+^-sensitive and K^+^-uptake deficient yeast was used to functionally characterise rice OsCCC1 as a K^+^ and Na^+^ permeable cotransporter (Chen *et al*., 2016). We used this approach and expressed VvCCC-YFP in yeast and stained cells with the FM4-64 dye, which initially stains the plasma membrane of live cells by intercalation, before being internalised (Fig. 4B). Here, the perimeter of yeast cells and specific intracellular compartments remained visible when stained with FM4-64 (Fig. 4B). However, VvCCC-YFP did not co-localise with any of the FM4-64 signal, indicating that this protein was absent at a plasma membrane or endocytic vesicles of yeast, which excluded this approach from using it in transport studies. However, as VvCCC is known to localise to the plasma membrane of *Xenopus* oocytes (Henderson *et al*., 2015), oocytes were used for transport studies.

### VvCCC displays KCC-like ^86^Rb flux dynamics in Xenopus oocytes

Payne (1997) calculated the thermodynamic driving forces for hKCC2 in neurons as a function of external K^+^. Similar calculations were performed for NKCC (DeFazio *et al*., 2000), who predicted the expected outcomes of NKCC and KCC flux experiments in *Xenopus* oocytes at different Cl^-^ internal concentrations ([Cl^−^]_i_). In *Xenopus* oocytes, CCC-mediated tracer uptake is typically investigated after pre-incubation in Cl^−^-free medium, which reduces [Cl^−^]_i_ to negligible levels (Gamba *et al*., 1994). We adopted the alternate approach and found out that at low [Cl^−^]_I_, the chemical driving force (Δµ) for NKCC transport was negative in standard ND96 solution (96 mM NaCl; 2 mM KCl), meaning that the driving force for coupled 1Na^+^:1K^+^:2Cl^−^ co-transport was inward (Fig. 5A, black line). By contrast, KCC operated much closer to the predicted flux reversal point (FRP), where net flux equalled zero (Fig. 5A, blue line). Cl^−^-depleted oocytes that expressed KCC would have a negative Δμ initially, but FRP would be reached when [Cl^−^]_i_ accumulated to 2.4 mM, compared to 46.7 mM for NKCC (Fig. 5A, X-axis intercepts). Thus, in oocytes with depleted of [Cl^−^]_i_, thermodynamics predicts that ^86^Rb uptake through a KCC would rapidly achieve equilibrium, while the NKCC protein would continue to mediate ^86^Rb influx for longer periods (Fig. 5A).

**Fig. 5:**
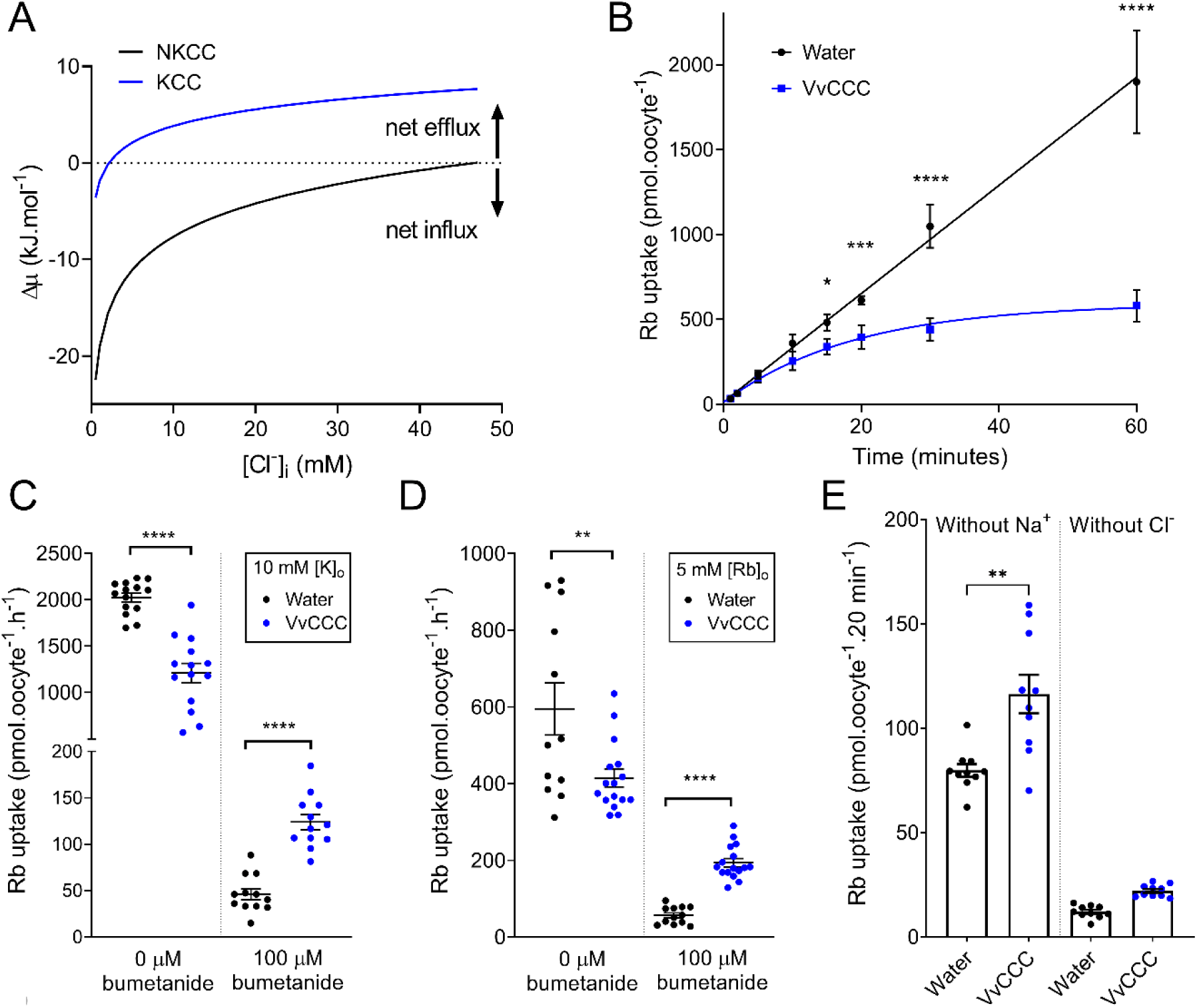
VvCCC-mediated ^86^Rb tracer fluxes display thermodynamic hallmarks of KCC cotransporters. (A) Predicted thermodynamic driving forces for KCC and NKCC co-transport into *Xenopus* oocytes as a function of [Cl^−^]_i_ in the ND96 solution with 106.6 mM [Cl^−^]_o_ 96 mM [Na^+^]_o_, and 2 mM [K^+^]_o_. Driving forces were calculated assuming 100 mM [K^+^]_i_ and 10 mM [Na^+^]_i_ at 25 °C using the following equations: 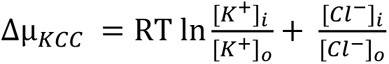 and 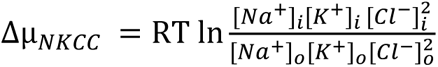 where R is the gas constant (8.314 kJ.mol^-1^.K^-1^) and T is the absolute temperature. Dashed line represents the flux reversal point (FRP). (B) Time course of ^86^Rb uptake in oocytes injected with VvCCC or water in standard ND96 solution. Oocytes were pre-incubated in Cl^−^-free ND96 overnight before measurements (Cl^−^ replaced with gluconate). Each data point is the mean ± Standard Deviation (SD, calculated in Microsoft Xcel 2019) of six oocytes (water) or eight oocytes (VvCCC). Asterisks represent significant differences at single time points (one-way ANOVA). (C– D) ^86^Rb uptake in oocytes injected with VvCCC or water after 1 hour in ND96 with 10 mM KCl (c) or 5 mM RbCl (D) in the absence or presence of 100 µM bumetanide. Bars indicate mean ± Standard Error of the Mean (SEM, calculated in Microsoft Xcel 2019). Asterisks represent significant difference between means (unpaired t-test). (E) ^86^Rb uptake in oocytes injected with VvCCC (blue) or water (black) after 20 minutes in ND96 without Na^+^ (left) or without Cl^−^ (right). Bars indicate mean ± SEM. Asterisks represent significant difference between means (unpaired t-test). For all panels, asterisks denote *P < 0.05 **P < 0.01 ***P = 0.001 ****P < 0.0001.

VvCCC flux dynamics in ND96 solution was compared to thermodynamic predictions using a one-hour time course of ^86^Rb uptake by Cl^−^-depleted oocytes. VvCCC-injected oocytes reached saturation with a half-time of 13.23 minutes determined from the one-phase association curve (R^2^ = 0.92) (Fig. 5B). By contrast, water-injected (control) oocytes accumulated significantly more ^86^Rb and the uptake displayed a linear relationship (R^2^ = 0.96) (Fig. 5B). These results are consistent with the thermodynamic predictions that VvCCC mediates KCC co-transport, and that NKCC co-transport occurs in water-injected control oocytes, which is mediated by their endogenous NKCC (Suvitayavat *et al*., 1994) (Fig. 5A). ^86^Rb uptake by control oocytes was almost completely inhibited by 100 µM of the NKCC blocker bumetanide, supporting that endogenous NKCC is responsible for ^86^Rb uptake in control oocytes (Fig. 3C; Fig. 3D). When [K^+^]_o_ was increased from 2 mM to 10 mM, ^86^Rb uptake by VvCCC-injected oocytes increased, while ^86^Rb uptake by water-injected controls remained unchanged (Fig. 3B; Fig. 3C; Supplementary Fig. S2). This agrees with a predicted shift in the FRP parameter for a KCC-mediated uptake mechanism by VvCCC, but lacking shift in FRP for the NKCC-mediated uptake mechanism, present in control oocytes, was observed (Supplementary Fig. S2). Replacing [K^+^]_o_ with 5 mM Rb^+^ led to similar outcomes but with a reduced total ^86^Rb uptake, which might indicate differences in VvCCC selectivity, or batch-to-batch expression differences (Fig. 5D). Unlike the water-injected control oocytes, ^86^Rb uptake by VvCCC-injected oocytes was not completely blocked by 100 µM bumetanide (Fig. 5C; Fig. 5D). This is consistent with previous findings of ^86^Rb uptake by VvCCC-injected oocytes (Henderson *et al*., 2015). Bumetanide affinity to KCCs is significantly lower than that of NKCCs. NKCCs have a typical half-maximal effective concentration (EC_50_) of ∼ 0.1 µM, while the typical EC_50_ value for KCCs is ∼ 100 µM bumetanide (Russell, 2000).

To determine whether VvCCC-mediated ^86^Rb uptake was Na^+^ coupled, Na^+^ was removed from the medium. Fluxes were carried out for 20 minutes, as VvCCC would have already reached saturation. In Na^+^-free ND96, VvCCC-injected oocytes accumulated more ^86^Rb than water-injected controls (Fig. 5E). When Cl^−^ was removed from the flux solution, neither VvCCC-injected nor the water-injected controls were capable to accumulate significant ^86^Rb, suggesting that ^86^Rb uptake is Cl^−^-coupled for both transport pathways (Fig. 5E). Collectively, these data indicate that VvCCC displays the K^+^-Cl^−^-like cotransporter activity.

### VvCCC elicits a Na^+^ conductance in Xenopus oocytes

Previously, the expression of plant CCCs in oocytes led to greater ^22^Na influx, and it was concluded that plant CCCs are NKCCs (Colmenero-Flores *et al*., 2007; Henderson *et al*., 2015). To explore this alternative, two-electrode voltage clamping (TEVC) was performed in the ND96 solution. Whole-cell currents of VvCCC-injected oocytes were greater than those of water-injected controls (Fig. 6A to Fig. 6C). Currents were inward at physiological V_m_, consistent with a cation influx (Fig. 6D). Resting V_m_ of VvCCC-expressing oocytes was depolarised compared to water-injected controls (Fig. 6E). This is expected for oocytes in 100 mM [Na^+^]_ext_ that express an electrogenic Na^+^ conductance, as the Nernst equation predicts V_m_ will move towards the equilibrium potential for Na^+^. To determine whether this conductance was mediated by VvCCC, oocytes were pre-incubated and clamped in 100 µM of furosemide, which is a potent KCC inhibitor. Conductance of VvCCC-injected oocytes was not significantly changed by furosemide treatment (Fig. 6F). Furosemide also had no effect on the conductance of control oocytes, which was significantly lower than of the VvCCC-injected oocytes in the presence or absence of the furosemide inhibitor (Fig. 6F). No Cl^−^ conductance was observed in this case, but the conductance did mediate inward K^+^ currents under the high external K^+^ (Supplementary Fig. S3). Taken together, these results indicate that an electrogenic Na^+^ (and K^+^) conductance is activated in VvCCC-injected oocytes and is independent of electroneutral VvCCC function. This conductance was formerly unidentified and would likely contribute towards the ^22^Na uptake that was observed in earlier studies (Sam Henderson, unpublished). Consistent with our findings, the expression of a mosquito (*Aedes aegypti*) CCC (aeCCC2) in *Xenopus* oocytes induced the Na^+^ and Li^+^ conductance of a similar amplitude to VvCCC that was not inhibited by loop diuretics (Kalsi *et al*., 2019). Cation conductance might be an inherent property of aeCCC2 and VvCCC. However, the induction of endogenous channels by high levels of heterologous membrane proteins in *Xenopus* oocytes was also reported (Tzounopoulos *et al*., 1995). Whether AtCCC also elicits a cation conductance in oocytes is uncertain, which represents another experimental target.

**Fig. 6:**
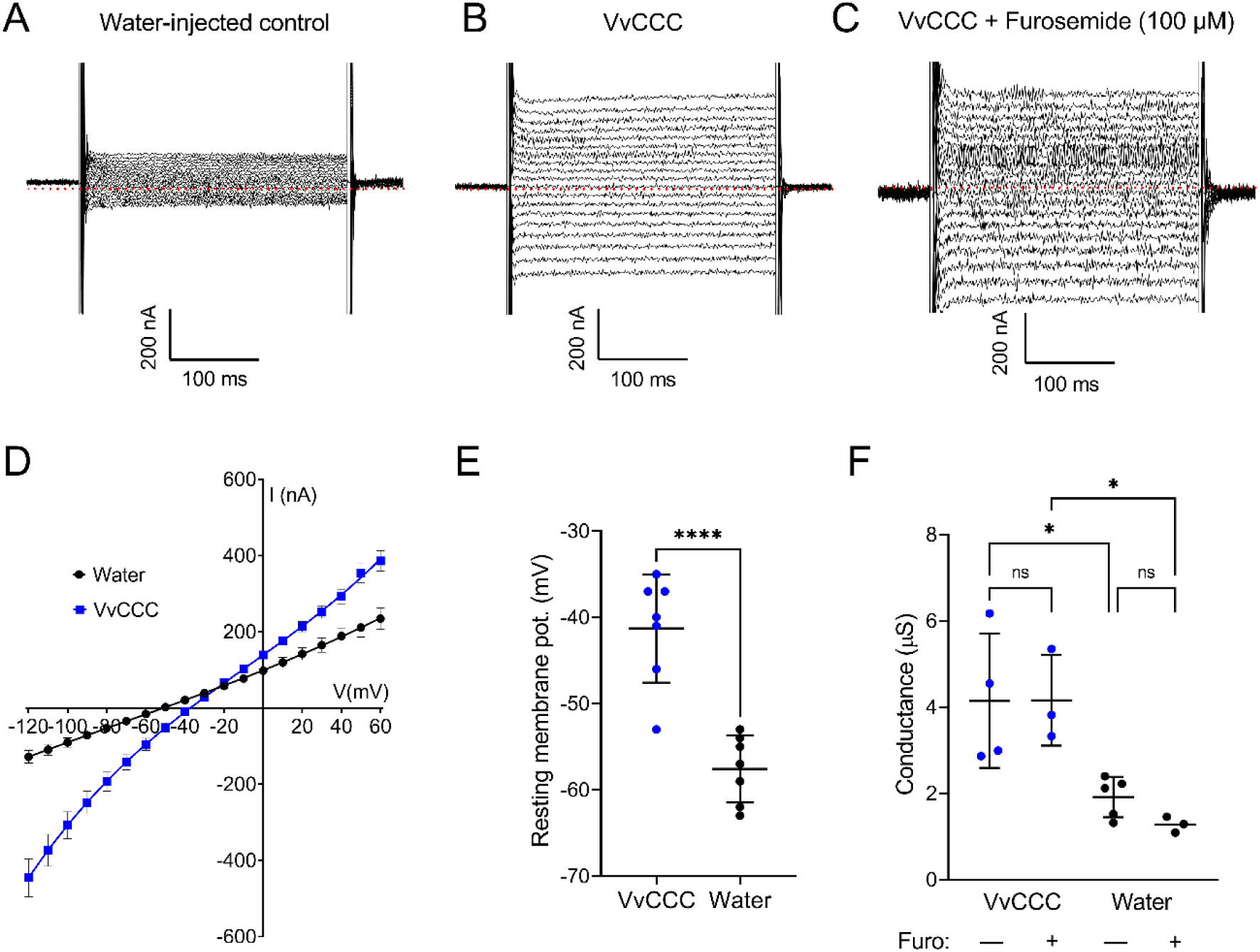
*VvCCC* expression induces an electrogenic cation conductance in *Xenopus* oocytes. (A – C) Whole-cell electrophysiology traces showing currents recorded from water-injected control oocytes (A), VvCCC-injected oocytes (B) and VvCCC-injected oocytes pre-incubated for 30 minutes and recorded in the presence of 100 µM furosemide (C). Dotted red lines denote zero current level. (D) Current-voltage relationships of oocytes injected with VvCCC (blue) or water (black) in ND96 solution. Data are the mean ± SEM of four oocytes. Curves were generated by fitting data with a polynomial function. (E) Resting membrane potential of oocytes injected with VvCCC (blue) or water (black) in the ND96 solution. Data indicate mean ± SEM. Asterisks denote statistically significant difference (unpaired t-test). (F) Whole cell conductance of oocytes injected with VvCCC (blue) or water (black) pre-incubated for 30 minutes and recorded in the presence or absence of 100 µM furosemide. Conductance was determined from the slope of the I/V curve close to the reversal potential. Asterisks denote statistically significant difference (one-way Anova with Tukey’s multiple comparison test). For all panels, asterisks denote *P < 0.05 ****P < 0.0001.

### How might a KCC-like protein function in plant cells?

The Golgi, TGN and EE are the part of the endocytic pathway in plants that is important for cellular trafficking and recycling to and from the plasma membrane (Sze and Chanroj, 2018). A progressive pH gradient (acidic lumen) is required for normal endosomal function, driven by the coordinated action of the proton (H^+^) pumping V-type ATPase along with a predicted Cl^−^/H^+^ exchanger (CLCd) and known K^+^/H^+^ exchangers (KEA, NHX5, NHX6, CHX17) to balance the charge. Arabidopsis AtCCC was proposed to complete the transport circuit in the TGN/EE through the coupled efflux of cations and anions out of the lumen towards to cytoplasm (McKay *et al*., 2022). Under non-stress conditions, [K^+^]_cyt_ is around 100 mM (Leigh and Wyn Jones, 1984) while [Cl^−^]_cyt_ is around 15 mM (Lorenzen *et al*., 2004; Saleh and Plieth, 2013). K^+^ and Cl^−^ concentrations within the TGN/EE lumen are unknown but likely to be greater than the cytoplasm since both ions are actively transported against their electrochemical gradients. This would support the CCC-mediated efflux mechanism from endosomes as suggested by McKay *et al*. (2022). The magnitude of CCC-mediated fluxes could be influenced by the combined activity of secondary active proton exchangers located at endo-membranes. CCC fluxes may also be influenced by plasma membrane K^+^ and Cl^−^ channels generating localised concentration gradients within cytosol, especially as the TGN/EE are mobile organelles.

If CCC activity is modulated, for example by phosphorylation and de-phosphorylation, the counter-ion concentration and thus the pH value within the TGN/EE lumen could be partially regulated. Multiple studies demonstrated that AtCCC is phosphorylated *in vivo* (Rayapuram *et al*., 2018; Reiland *et al*., 2009). Interacting partners of plant CCC proteins are currently unknown, however a Mitogen Activated Protein Kinase recognition sequence was identified at the N-terminal region of AtCCC (Sorensson *et al*., 2012). AtCCC and its co-localisation partner AtCLCd could both act as the negative regulators of pathogen-associated molecular pattern (PAMP)-triggered immunity (Guo *et al*., 2014; Han *et al*., 2020), which suggests a possible new signalling pathway that CCC proteins could be involved in.

In conclusion, this study demonstrates that plant CCCs, such as VvCCC could function as a K^+^-Cl^−^ symporters. These conclusions are supported by phylogenetic and structural predictions and experimental data presented in this work, which could pave the way for future correlative studies of plant CCCs.

## Supplementary data

**Supplementary Table S1:** Sequence identities/similarities between VvCCC or AtCCC, and DrNKCC1, hKCC1 and hKCC3 transport proteins, and protein modelling evaluation parameters of full-length (FL) and TMD proteins.

**Supplementary Fig. S1:** Comparison of VvCCC models using DrNKCC1 and hKCC3 cation-chloride cotransporter as templates.

(A) Cartoon representations of TMD domains (chains B) of DrNKCC1 (yellow), and VvCCC (cyan) based on the TMD template of DrNKCC1, illustrating poses of K^+^ and two Cl^−^ ions (cpk spheres) in pores.

(B) Cartoon representations of TMDs (chains B) of the cryo-EM hKCC3 structure (light grey), and the VvCCC (dark grey) models based on the TMD template of hKCC3, illustrating poses of K^+^ and two Cl^−^ ions (cpk spheres) in pores.

(C) Superpositions of TMD domains (chains B) of the DrNKCC1 (yellow) and hKCC3 (grey) structures, the VvCCC model based on the TMD templates of DrNKCC1 (cyan) or hKCC3 (dark grey). Respective rmsd values of the superposed structures are 1.5 Å (472 and 544 residues), and 1.6 Å (460 and 472 residues).

**Supplementary Fig. S2:** The driving force for VvCCC-mediated ^86^Rb uptake is positively correlated with external [K^+^].

(A) Predicted thermodynamic driving forces for KCC and NKCC cotransport into *Xenopus* oocytes as a function of [Cl^−^]_i_ in ND96 solution with 2 mM [K^+^]_o_ or 10 mM [K^+^]_o_. Calculations were performed as described in Materials and methods. Flux reversal point is indicated by the intercept with the horizontal dotted line. Note that changes in [K^+^]_o_ alter the predicted FRP for a KCC mechanism (blue and red lines), but not for an NKCC mechanism (black line).

(B) ^86^Rb uptake in oocytes injected with VvCCC (blue circles) or water (black circles) after 1 hour in ND96 with either 2 mM KCl (left) or 10 mM KCl (right). Note that increasing [K^+^]_o_ only changed the magnitude of ^86^Rb uptake by VvCCC-injected oocytes. Data in (B) are also presented in Fig. 3B and Fig. 3C.

**Supplementary Fig. S3:** Selectivity of VvCCC-injected oocyte currents to K^+^, Na^+^ and Cl^-^.

Current-voltage relationships of oocytes injected with VvCCC at the indicated membrane potential (V) while bathed in 96 mM Na^+^ (ND96, blue), or the same solution where 96 mM NaCl and 2 mM KCl was replaced with 98 mM KCl (red), 98 mM NMDG-Cl (purple), or the standard ND96. Data are the mean ± Standard Error of the Mean (calculated in Microsoft Xcel 2019) of three oocytes after subtraction of the currents from a single water-injected control oocyte.

## Acknowledgements

The University of Adelaide School of Biomedicine, and the School of Agriculture Food and Wine, provided equipment and resources. Pak Chin How are thanked for providing the assistance with HEK293 cells and transfections. Andrea Yool is thanked for providing access to equipment and resources. Steve Tyerman, Andrew Moorhouse, and Matthew Gilliham are thanked for useful discussions. Matthew Gilliham is also thanked for providing the expression vector for oocyte experiments through CE140100008. The pCDNA3.2 vector was kindly provided by Sheryl Shoubridge. Wendy Sullivan is thanked for performing *Xenopus* oocyte surgeries. Adelaide Microscopy is thanked for assistance with confocal microscopy.

## Author Contributions

SWH performed experiments, analysed data, and wrote the manuscript. MH performed structural modelling and wrote the manuscript. SN assisted with HEK293 cells and transfections.

